# Allocating Resources across Time in a Cumulative Allocation Game

**DOI:** 10.64898/2026.10.29.762832

**Authors:** Yokyung Esther Wi, Keiji Ota, Tessa Dekker, Laurence T. Maloney

## Abstract

We investigate an alternative to decision under risk where players begin with an endowment of wealth, and their goal is to increase this wealth as rapidly as possible by repeatedly investing part of it. In such a Cumulative Allocation Game (CAG), expected wealth can grow exponentially but – if wealth ever goes to zero – the game is over and the participant wins nothing (“ruin”). CAGs can be used to model a wide range of biological and economic activity from casino gambling to animal foraging to stock market investment (Kelly Criterion). However, there is little previous research concerning human performance in CAGs and how it relates to human performance in similar decision-under-risk tasks. In two experiments we asked participants to complete four CAGs designed to allow us to test a series of hypotheses about human performance and compare it to normative play maximizing their log rate of growth. In the main experimental conditions, participants’ performance was initially close to normative but their proportion bet decreased by almost a factor of two over the course of the experiment. This reduction in proportion-bet decreased their expected rate of growth by roughly 25% -- a substantial deviation from optimal. *Why would participants systematically abandon a normative strategy?* We conjecture that a possible reason is loss aversion, well documented in decision under risk. We conjecture that while loss aversion in decision under risk is sub-optimal, there is an intrinsic asymmetry between losses and gains in CAGs that could explain the loss aversion we observe.

## Introduction

Over the course of a day, we allocate our limited resources - time, energy, money – to improve our standing now and in the future. We risk part of what we have to gain more. In allocating our resources between different activities, we should consider not only our immediate gains and losses but also how our allocation strategy affects our future gains and losses. If we choose well in a Cumulative Allocation Game (CAG), our resources can grow exponentially; if we choose badly, we may be left with little or nothing to allocate. Running out of time, energy or money - in economic terms, “ruin” (Ramsey, 1995; Oprea, 2014) – is potentially disastrous for any organism.

When a CAG is a “favorable game” in Breiman’s terminology (Breiman, 1960, 1961), the focus is on maximizing the log rate of increase of player’s “wealth” while avoiding ruin in a game that can last an indefinite number of turns. There is a substantial mathematical literature on favorable games (Kelly, 1956; Thorp, 1969) and their application to the stock market (MacLean, Thorp & Ziemba, 2011), to casino games, notably blackjack (Thorp, 1969) and to horse racing. Poundstone (2005) provides an elegant history of the people and ideas involved. Staddon (1980) collects research on allocation games drawn from many fields including economics and ethology. Other representative studies concern allocation optimization in ethology and in population genetics (Baddeley, Franks & Hunt, 2019; Donaldson-Matasci, Lachmann & Bergstrom, 2008; Wei & Wang, 2024; Tal & Tran, 2020). Okabe (2022), for example, reformulates natural selection as a CAG.

There is little experimental research on human performance in CAGs and little attempt to connect human behavior and its frailties in decision under risk (Kahneman & Tversky, 1979; Tversky & Kahneman, 1992) to behavior in CAGs. This is surprising since a CAG is evidently a plausible model of both biological foraging and economic activity. We examined how human players actually play a simple CAG and compare their performance to normative play.

### The Cumulative Allocation Game

The single-player game we consider is played in turns. On the *n*^th^ turn of the game the player has a total wealth *W*_*n*_ expressed in *tokens*. Initially the player is given a total wealth of *W*_*1*_ *= 30* tokens.

Figure 1 illustrates one turn of play. At the beginning of the *n*^th^ turn the player is offered a *wager* with odds, *O*_*n*_. The odds are equally likely to be 2:1,3:1 or 4:1. The player is invited to bet some, all or none of his total wealth *W*_*n*_ with the understanding that, with probability 0.5, he will receive nothing back (he has lost his bet and the tokens he bet) and, with probability 0.5, he will receive his bet *B*_*n*_ back multiplied by the odds *O*_*n*_. Figure 1 illustrates a worked example.

**Figure 1:**
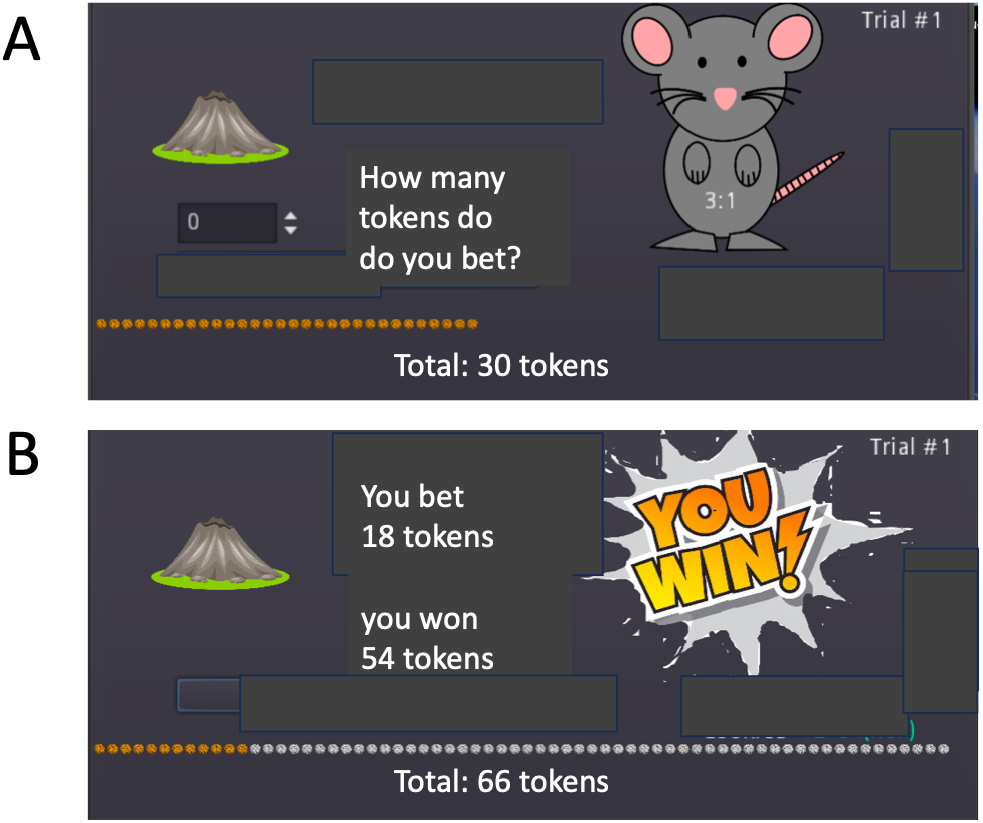
A single turn of the cumulative allocation game (CAG). On each turn the player sees a computer display that contains the information needed to play the game. We illustrate possible play on the first turn at two points during the turn. **A. The wager**. Initially the player has *W*_*1*_ *=* 30 tokens and sees a numerical summary of his current wealth, an array of 30 tokens. The player is also shown the odds *O*_1_ on the first turn which can be 2:1, 3:1, or 4:1 with equal probability. In our example, *O*_*1*_ *=* 3:1, displayed on the cartoon mouse. The player bets any amount from 0 up to his current wealth (30 tokens) and he wins or loses tokens depending on the outcome. Suppose he chooses to bet 18 tokens. He now has 12 tokens remaining (the amount he did not bet). **B. The outcome**. With probability 0.5 the player receives nothing back (he loses the 18 tokens he bet) and, with probability 0.5, he receives *O*_1_ *B*_1_ (his bet multiplied by the odds). The figure shows the display as if the player has won and now has *W*_*2*_ = 12 + 3 × 18 = 66 tokens at the start of the second turn. Had he lost he would have had only 12 tokens, the tokens not bet. At the end of each turn a cartoon volcano may erupt with probability 1/30, ending the game (see Stopping Rules below). On the second trial, he can now bet any portion of his current wealth, 66, with odds *O*_*2*_. The actual graphical displays were designed to be used with children as well as adults in later experiments. Consequently, the odds are presented as a friendly mouse, the tokens as cookies, and the end of the game as a volcanic eruption.

On each turn of the game the player’s total wealth can increase or decrease. If the player’s wealth is reduced to only a few tokens he will recover his losses only with difficulty. Growth, though exponential, can be slow. If the player’s total wealth ever falls to precisely 0 then the player is ruined, the game is over, and the player receives no bonus for performance. If the player avoids ruin and successfully reaches the end of the game (defined by a stopping rule described below), he is paid a monetary bonus proportional to his total wealth in tokens at the end of the game. We compare human performance to a normative strategy (Kelly, 1956; see also Thorp, 1969) that maximizes the rate of growth of expected log wealth, E[ logW ].

The expected value *EV*_*n*_ of a bet *B*_*n*_ with odds *O*_*n*_ (the expected value or loss to the player on a single turn) is

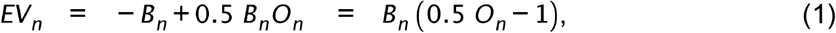

proportional to the size of the bet). If *O*_*n*_ *<* 2 the expected value of the bet is negative and the normative player should refuse to bet by setting *B*_*n*_ = 0. If *O*_*n*_ *=* 2 the expected value is 0 and the game is equivalent to flipping a fair coin and receiving −*B*_*n*_ or *B*_*n*_ depending on the outcome of the coin toss. The expected value is 0 and we would expect that the player with any degree of loss aversion (Markowitz, 1952; Kahneman & Tversky, 1979) or risk aversion (Bernoulli, 1738/1954; Luce, 2000) will not bet when *O*_*n*_ *=* 2. We include such trials precisely to determine how players will react to gambles with expected value 0 interleaved among gambles with positive expected value. Will they (correctly) refuse to bet?

If *O*_*n*_ > 2 the expected value *EV*_*n*_ on each turn is positive and, as just noted, proportional to *B*_*n*_. Since the odds are in the Player’s favor, the game is a “favorable game” in Breiman’s terminology (Breiman, 1960, 1961 and the player’s expected wealth can grow exponentially (Kelly, 1956; Bellman & Kalaba, 1957; Breiman, 1960, 1961; Thorp, 1969).

**Table 1:**
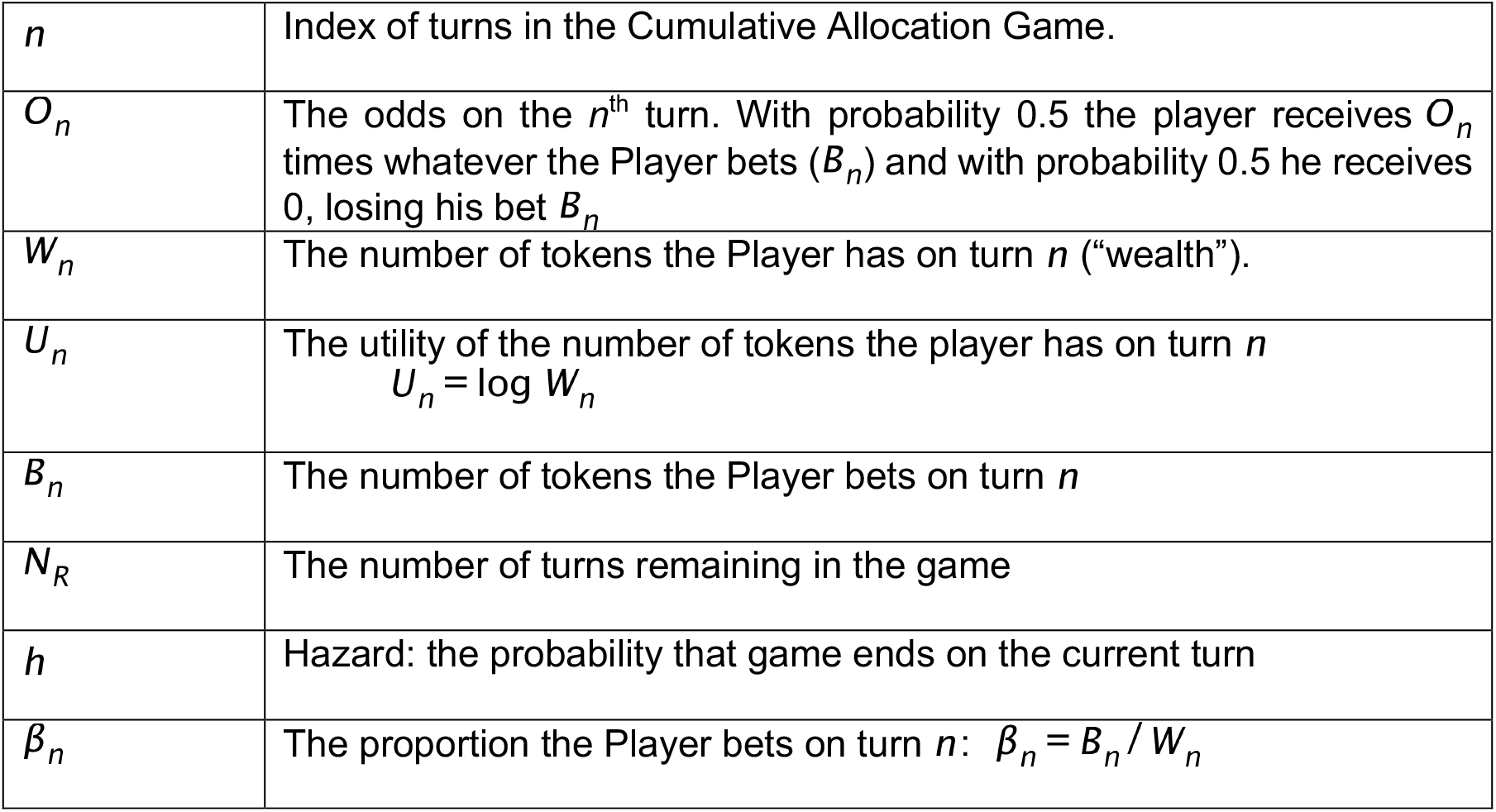
Notation. When any of these are followed by an asterisk (e.g. p*) the normative value of the variable is intended.

|  |  |
|---|---|
| *n* | Index of turns in the Cumulative Allocation Game. |
| *O<sub>n</sub>* | The odds on the *n*<sup>th</sup> turn. With probability 0.5 the player receives *O<sub>n</sub>* times whatever the Player bets (*B<sub>n</sub>*) and with probability 0.5 he receives 0, losing his bet *B<sub>n</sub>* |
| *W<sub>n</sub>* | The number of tokens the Player has on turn *n* (“wealth”). |
| *U<sub>n</sub>* | The utility of the number of tokens the player has on turn *n*<br>*U<sub>n</sub>* = log *W<sub>n</sub>* |
| *B<sub>n</sub>* | The number of tokens the Player bets on turn *n* |
| *N<sub>R</sub>* | The number of turns remaining in the game |
| *h* | Hazard: the probability that game ends on the current turn |
| *β<sub>n</sub>* | The proportion the Player bets on turn *n*: *β<sub>n</sub>* = *B<sub>n</sub>* / *W<sub>n</sub>* |

### Stopping Rules

In playing a CAG, we should consider how the game might end (the stopping rule). The game could go on indefinitely or end after a fixed number of turns known to the player. When only a few turns are left, how should players alter their strategies if at all? A possible stopping rule is,

#### The known duration stopping rule

The player is told exactly how many trials *N* the game will last (assuming the player avoids ruin). When the stopping rule is the known duration stopping rule we include a countdown timer in the game display specifying how many turns remain, *N*_*R*_.

How should the player react as he approaches the end of the game?

Suppose he knows that the current turn is the last turn of the game: *N*_*R*_ = 1. The normative strategy maximizing expected wealth is perhaps surprising. If as we assume, the player wishes to maximize expected wealth then, when *O*_*n*_ *=* 3 or *O*_*n*_ *= 4* he should bet *all* of his *W*_*n*_ tokens. We refer to this strategy as the *“bet-it-all” strategy*. When *O*_*n*_ *=* 3 or *O*_*n*_ = 4 the expected value is an increasing function of the bet *B*_*n*_ (Eq. 2) and is maximized when the bet *B*_*n*_ is maximized as it is with the “bet-it-all” strategy. When *O*_*n*_ = 2 the expected value is 0 for any strategy including the “bet-it-all” strategy.

The last turn, however, is not special. If the player knows that the game will end after two turns and wishes to maximize expected value, he should still “bet-it-all” on each of the two turns remaining (Kelly, 1956; Ferguson, 1965). He should first maximize his winnings on the second last turn and then maximize his winnings on the last turn by betting all his accumulated wealth including his winnings on the previous turn. This strategy maximizes his expected winnings across the last two turns (even taking into account the possibility of ruin on either turn). It can be generalized: if there are a fixed and known number of turns remaining, he should “bet-it-all” on each turn.

Even though the bet-it-all strategy maximizes expected value it is implausible (to us) that any human player would follow such a strategy. The probability of ruin on or before the *N*^*th*^ *turn* is 1 − 2^−*N*^, rapidly converging to 1 as the number of turns remaining (*N* ) increases. The player following such a strategy for a substantial number of turns will almost certainly be ruined but − if not ruined − fantastically wealthy.

In Experiment 2, we will compare what human players do when confronted with this fixed duration stopping rule and an alternative stopping rule − constant hazard − designed so that the player can never anticipate when the game is about to end.

#### The constant hazard stopping rule

With the constant hazard stopping rule, players are told that the game could terminate without warning at the end of any turn with constant probability (*hazard*) *h* = 1/30. The “hazard events” at the end of each turn are independent, identically distributed, and the duration of the game *D* with a constant hazard stopping rule is a geometric random variable (Evans, Hastings & Peacock, 2000). The game with constant hazard is “memoryless” in the sense that no matter how long the game has gone on, the conditional probability that it will end after one more trial (the *hazard rate h*) is - by design - constant. Even after many turns the player has no more reason to expect that the game will end in one more turn (or two or three …) than he did at the start of the game.

### Normative Performance in Constant Hazard CAGs

What is the normative strategy over the course of the game when the stopping rule is constant hazard? At one extreme, the player could − on each trial − refuse to bet anything. He would retain whatever he had at the start of the trial. At the other extreme he could bet everything he has on each trial with an ever-increasing probability of ruin. An intermediate strategy is to bet some fixed proportion *β* of his current wealth *W*_*n*_ on each trial. For example, on every turn a player might bet half of his current *W*_*n*_, reserving the remainder as a buffer against ruin. Kelly (1956; Breiman, 1961) showed that the choice of *β*_*n*_ (the proportion to bet on the n^th^ turn) that maximizes the rate of growth of E[ logW ] is^1^

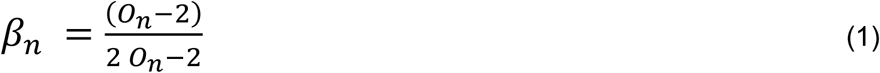

The normative proportion to bet depends only on the odds. Figure 2 is a plot of normative proportion bet vs. odds (Eq. 1).

**Figure 2:**
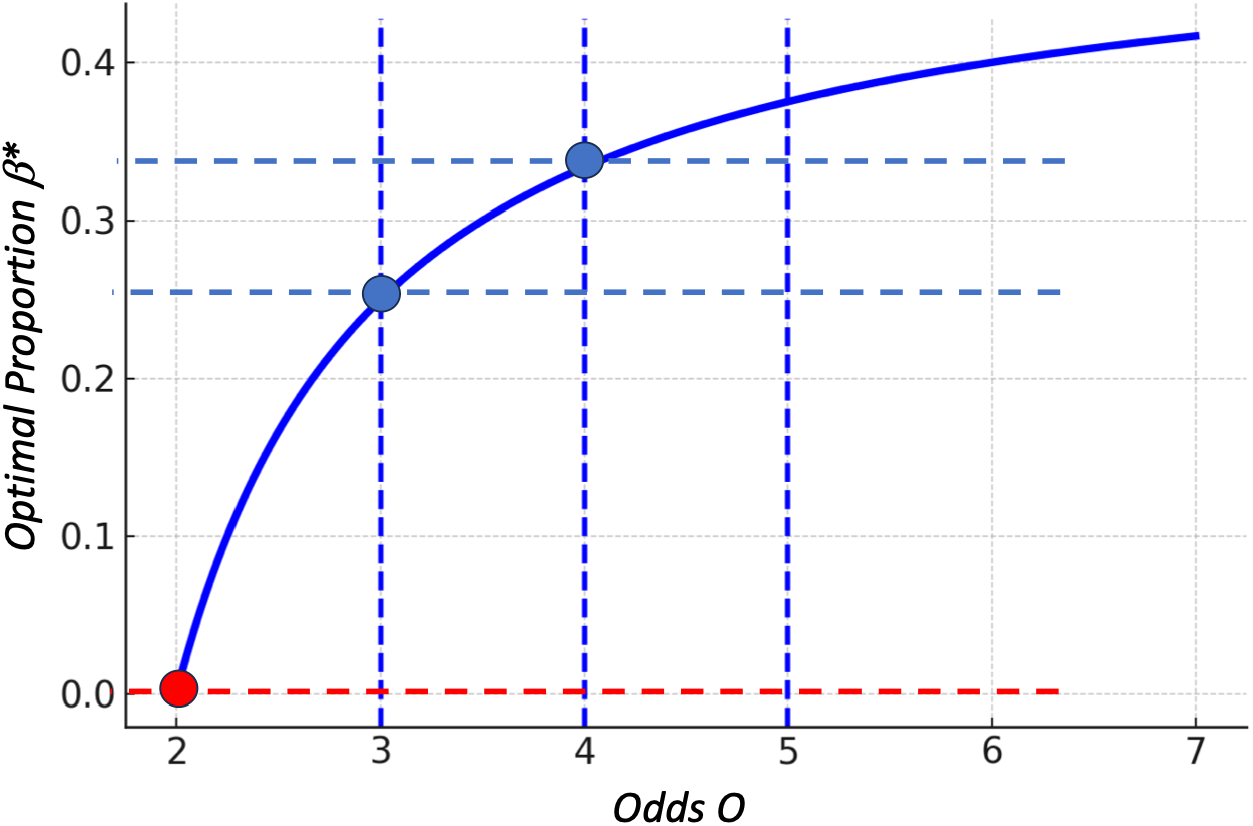
Normative proportion *β*\* versus odds O. The experimental conditions are marked by solid disks. The red disk corresponds to odds 2:1 where the optimal strategy is not to bet (*β*\* = 0).

#### Properties of Normative Play

To simplify analysis, we derive a series of properties of normative play assuming the player’s goal is to maximize the expected rate of growth of E[ logW ]. We do not assume that actual players will adhere to any of these properties, but it is useful to compare human performance to ideal (Geisler, 1989).

##### Property 1

**[Proportionality] when the stopping rule is constant hazard and** *O*_*n*_ *>* 2 **then normative strategy maximizing the expected rate of increase in wealth is to bet a fixed proportion** *β* > 0 **of current wealth** *W*_*n*_. **When** *O*_*n*_ ≤ 2, **the normative strategy is to bet** *β = 0* . The proportion bet does not depend on *W*_*n*_ or *n*. It depends only on *O*_*n*_ the odds offered on the current turn *n*.

The player may satisfy Property 1 while picking a fixed value of *β* that is too big or too small to maximize the expected rate of increase. If the player picks the normative value (Eq. 1, maximizing the expected rate of increase for each *O*_*n*_ ) he satisfies an additional property.

##### Property 2

**[Optimal-Bet] When the stopping rule is constant hazard and** *O*_*n*_ *>* 2 **bet the proportion** *β***\* that maximizes the expected rate of growth of current wealth** *W*_*n*_ . We calculated these normative values using Eq. 1 (plotted in Figure 2) for each of our experimental conditions. For *O*_*n*_ = 3 the normative proportion was 0.25 and for *O*_*n*_ = 4 the normative proportion was 0.33. If Property 1 fails, then Property 2 cannot hold.

We add one more property of normative play that applies only to games with fixed duration stopping rules. In such games, the normative amount *B*_*n*_ maximizing expected value, on any trial is *B*_*n*_ = *W*_*n*_: the normative strategy is to “bet-it-all” on every trial.

##### Property 3

**[Bet-It-All] When the stopping rule is fixed duration and** *O*_*n*_ *>* 2, **the normative strategy maximizing expected value is to always bet-it-all ( *β* = 1 )**.

## Methods & Material

We report two experiments. Each experiment consisted of two plays of the CAG game referred to as *Game 1* and *Game 2* (Figure 3). A player participated in only one of the experiments, completing both games in the assigned experiment. All games lasted exactly 36 trials but in the two games in Experiment 1 and the first game of Experiment 2 the player did not know this.

**Figure 3:**
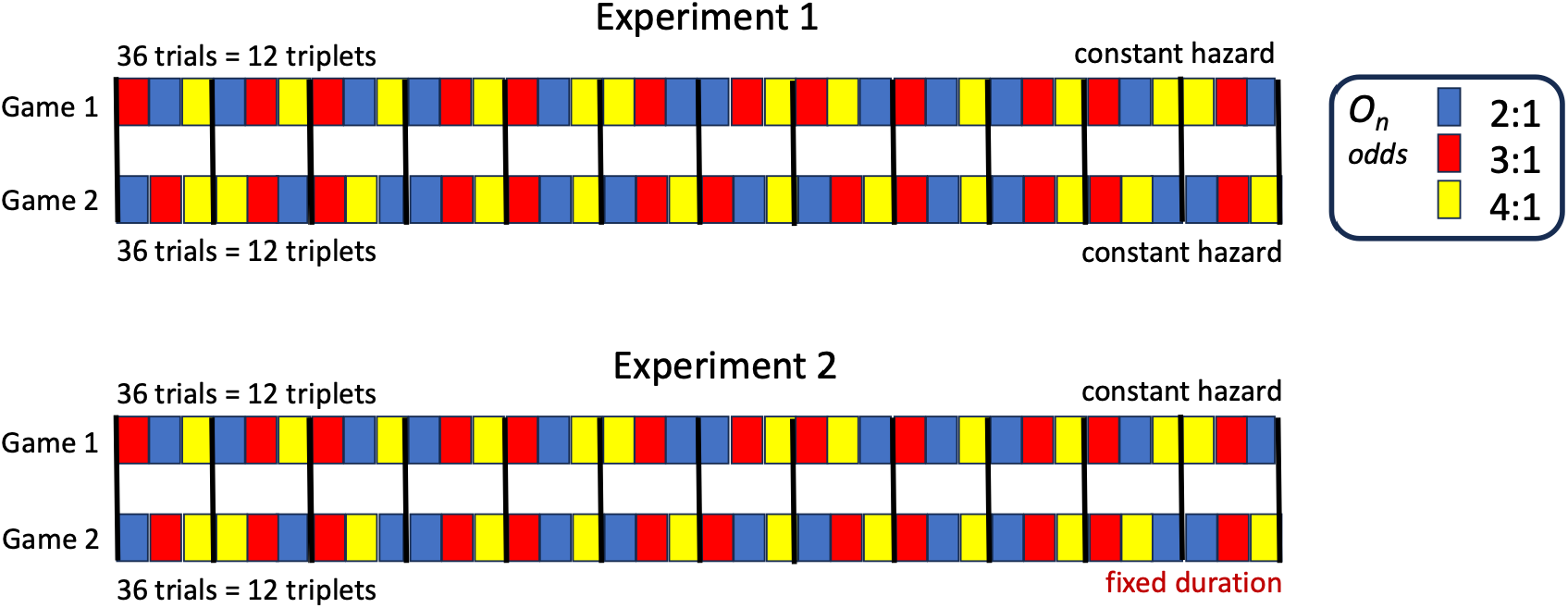
Structure of the experiments. Each player was randomly assigned to either Experiment 1 or to Experiment 2 but not both. Each experiment consisted of two games, each consisting of 36 trials of the CAG. For convenience, we partitioned the 36 trials in a game into 12 successive groups of three trials called *triplets*. Each of the twelve triplets in a game consisted of one each of the three gambles *O*_*n*_ *=* 2, 3 *or* 4 in randomized order. Each triplet contains one trial for each of the three possible odds. All possible triplets of 2, 3 and 4 occurred equally often. *The randomizations shown are not the randomizations actually employed*. In both games of Experiment 1 and in the first game of the second experiment players were told that the stopping rule was constant hazard. In the second game of the second experiment players were told that the play would last exactly 36 trials and a countdown clock displayed the number of turns remaining. In actuality each of the four games across two experiments lasted exactly 36 trials

In Experiment 1, the player was told that stopping rules for both games was constant hazard (the concept of constant hazard was explained and tested in training). The two games in Experiment 1 were objectively identical except for the randomized order of trials. We can therefore look for evidence of learning or other effects of experience by comparing the player’s performance in the two games in Experiment 1.

In Experiment 2 we examined the effect of the stopping rule. Players were given constant hazard instructions for Game 1 but were told that Game 2 would end after 36 trials (fixed duration) if the player avoided ruin. The experimental design is schematized in Figure 3.

### Players

Twenty players participated in Experiment 1 and twenty more players participated in Experiment 2. One player in Experiment 1 was replaced because they did not finish the experiment. Their data were not analyzed. Game 1 in Experiment 1 and Game 1 in Experiment 2 are identical, allowing us to test the replicability of our results.

All players gave informed consent before the beginning of the experiment. All were unaware of the purpose of the experiment. All were paid $12 per hour for participation plus a bonus proportional to their total wealth at the end of the experiment (each token was valued as $0.01). The one player who did not complete Experiment 1 was paid $12/hour for their time in the experiment. All experiments were approved by the Institutional Review Board at New York University (IRB-FY2022-6764).

### Apparatus

Stimuli and training material were presented on an ASUS display screen controlled by a Mac Minicomputer. The experimental program was written in GODOT4. Players were seated approximately 70 cm from the screen and made all responses by pressing buttons on a computer keyboard and/or moving a computer mouse.

### Procedure

#### Training

The players first completed a detailed Powerpoint® tutorial which explained the rules of the game. They were quizzed during training to verify that they understood the rules, in particular, the stopping rule with constant hazard.

### Design: Experiment 1

The first experiment consisted of two games, each comprising 36 turns (Figure 3). We compared the two games to detect any effect of experience. On each trial the player would see the current total wealth as a pile of *W*_*n*_ tokens (Figure 1A). If the number of tokens at the end of the turn ever reached 0, the experiment ended, and the player received no bonus (“ruin”). If the player did not reach ruin, each of the two games in reality had fixed durations of 36 turns each but players were told that the duration of the game was variable and that the game followed the constant hazard stopping rule.

At the end of each turn the controlling computer program would appear to “check” whether the game had come to an end with hazard probability *h =* 1/30 or would continue. When the first game ended after 36 turns, the player had no reason to suspect that its duration was not produced by a process of constant hazard. After the second game also ended at 36 trials the player might suspect that something was awry, but by then the experiment was over for that player. The deception and the reason for the deception were explained to the player following the experiment.

### Design: Experiment 2

Experiment 2 was identical in design to Experiment 1 except that in the second game the player was told that the game would end after precisely 36 turns (the fixed duration stopping rule). In the second game (with the fixed duration stopping rule) a countdown timer appeared on the screen counting the turns remaining. This design allowed us to assess the effect — if any — of the stopping rule.

## Results

We compared human performance to ideal performance maximizing the rate of growth of E[ logW ]. We summarize the main results of Experiment 1 in the following grey text box followed by full details of the statistical analysis for Experiment 1.

**Summary: Experiment 1 [Proportionality, Optimal-Bet]**

A. In all but one condition players failed Proportionality. Their proportion bet decreased over the course of Game 1 and Game 2 by roughly a factor of 2.
B. In both games when *O*_*n*_ = 3 or *O*_*n*_ = 4 players *initial* bets were indistinguishable from Optimal-Bet.
C. Game 2 is just a repetition of Game 1 with the same player. Comparing the outcomes of Game 1 and Game 2 we found no evidence of learning or other effects of experience.
D. When *O*_*n*_ = 2 in Game 1 and Game 2 the normative betting strategy was to bet 0. Players bet significantly more than 0 in both Game 1 and Game 2 although their bets in Game 2 showed roughly the same decrease toward zero as was found for *O*_*n*_ = 3 and *O*_*n*_ = 4 in Game 1 but not Game 2.

### Details of the analysis: Experiment 1: Proportionality, Optimal-Bet

We tested proportionality for each value of odds *O*_*n*_. In each 36-trial game the Player will encounter each of the odds *O*_*n*_ *=* 2,3,4 twelve times, once in each triplet of trials. The normative player maximizing E[ logW ] would bet the same proportion of his current wealth *β*_*n*_ on the same odds.

In Figure 4 we plot the proportion bet *β*_*n*_ for each player versus the trial number *n* separately for each value of odds *O*_*n*_ in both games of Experiment 1. There are three panels ABC for the three odds of game 1 and three corresponding panels DEF for game 2. If proportionality holds, we expect the plots (red lines marking the mean proportion bet versus triplet) to have no trend.

**Figure 4:**
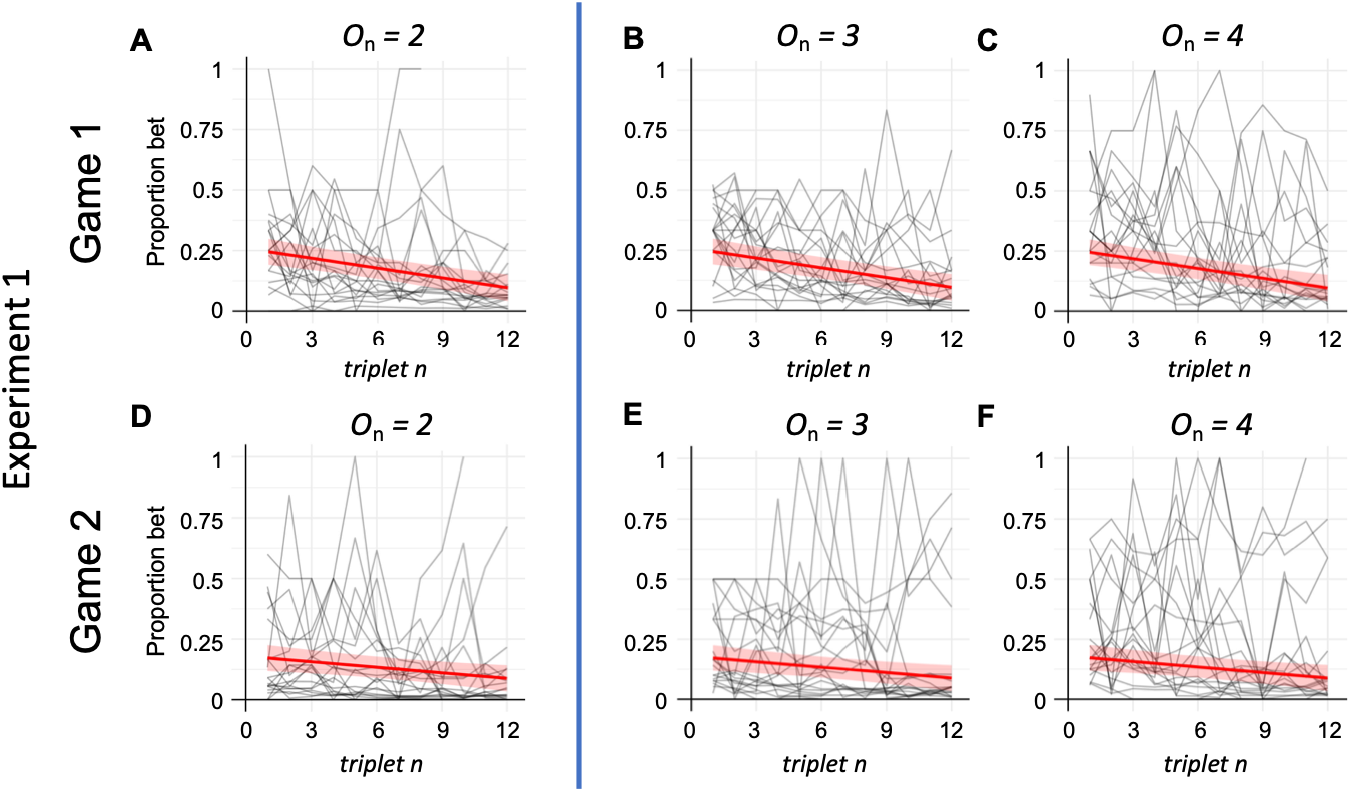
Proportion Bet in Game 1 and Game 2 of Experiment 1. Game 2 is a repetition of Game 1 (with a different randomization) allowing us to assess whether Players’ performances changed with experience in the task. **ABC Plots of proportion bet versus turn *n* for respectively for Game 1. DEF Plots of the same for Game 2**. The red lines are fitted regression lines. The Player maximizing growth of E[ logW ] would show no trend across trials with increasing trial number.

#### Experiment 1

*O*_*n*_ **= 3 and for** *O*_*n*_ **= 4: [Optimal-Bet]** We first report the results for *O*_*n*_ 3 and for *O*_*n*_ = 4 where following the normative strategy should lead to exponential growth and for O., = 2 where the normative strategy is never to bet.

The means of proportion bet and 95% confidence intervals in Figure 4BCEF were obtained using the following linear regression equation with a random intercept for each participant:

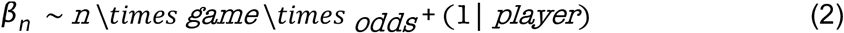

where *n* is trial number, game an indicator variable (0 for game 1, 1 for game 2) and *odds* is also an indicator variable (0 for *O*_*n*_ = 3 and 1 for O_n_ = 4). We performed a separate analysis on *0*_*n*_ = 2 with the equation below:

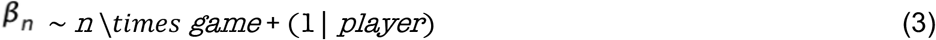

This model allows us to test for proportionality by modelling if the proportion bet over time stays constant -> slope=0), sensitivity to different odds by contrasting O3 and O4, and learning effects by comparing slopes across game 1 and 2. The estimated means and 95% confidence intervals are plotted in Figure 4AD. Given the mixed-effects model, we obtained the estimated marginal trends and the estimated marginal means when the trial was fixed at the first trial or the last trial. The player’s proportion bet *β*_*n*_ on the first trial of game 1 was not significantly different from the normative proportion for both *O*_*n*_ = 3 *(β*_*n*_ = 0.287, 95% CI = [0.211, 0.364]) and for *O*_*n*_ = 4 *(β*_*n*_ = 0.337, 95% CI = [0.261, 0.414]). However, we found significant downward trends in the slopes of regression when the odds of Game 1 were *O*_*n*_ = 3 (slope = -0.014, 95% [CI = [-0.021, -0.007]) and *O*_*n*_ = 4 (Figure 4C; slope = - 0.015, 95% CI = [-0.021, -0.008]). The estimated proportion-bet on the last trial of game 1 was significantly less than the normative proportion for both *O*_*n*_ = 3 *(β*_*n*_ = 0.133, 95% CI [0.056, 0.211]) and for *O*_*n*_ = 4 *(β*_*n*_ = 0.178, 95% CI = [0.101, 0.256]). Players initially bet proportions that were not significantly different from normative but reduced the proportion bet over the 12 triplets in a game. We replicated this failure in Game 2 (panels 4B vs. 4E, panels 4C vs. 4F). We discuss this pattern of results further in conjunction with Figure 5.

**Figure 5:**
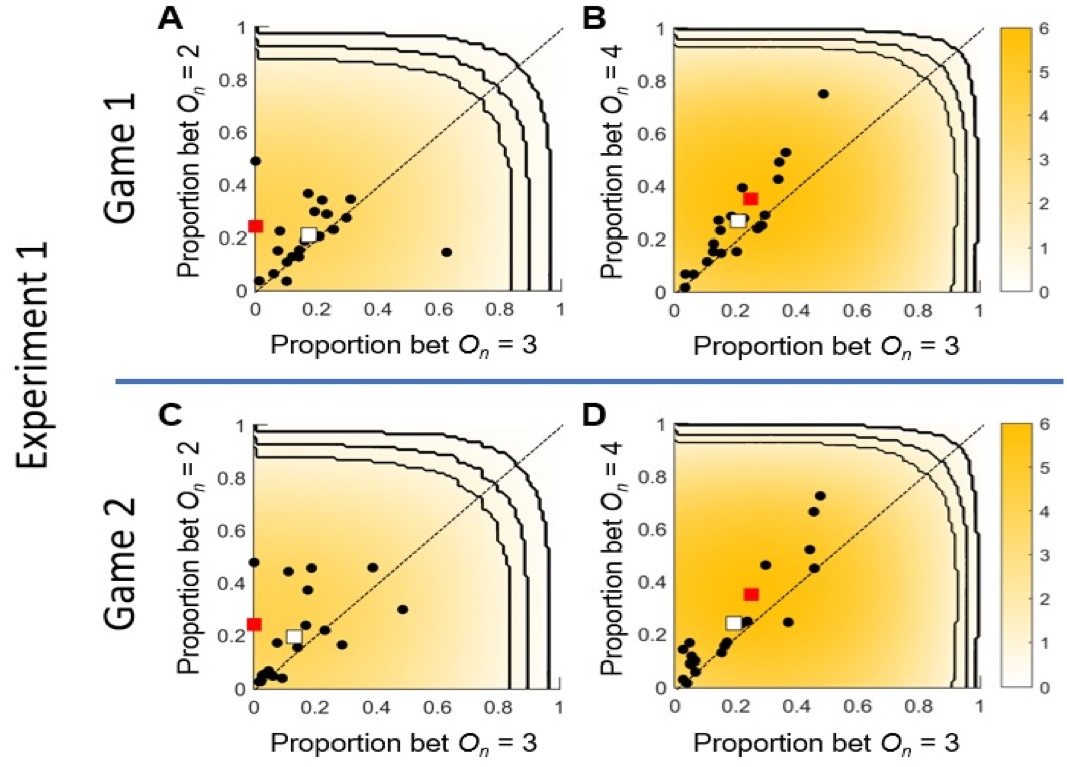
Experiment 1. Game 1. The mean proportion bet. A. Game 1. proportion bet O_n_ = 2 vs proportion bet O_n_ = 3. The normative proportion bet is plotted as a red square. Results for individual are players plotted as black dots. **The mean proportions bet across players re plotted as a white square. B. Game 1. proportion bet O**_**n**_ **= 4 vs proportion bet O**_**n**_ **= 3. C, D**. Game 2. same format as A, B respectively but for Game 2.

The results of Game 2 of *O*_*n*_ = 4 were not significantly different from those of Game 1. We found a significant downward trend in the slope of regression (Figure 4F; slope = -0.011,95% CI = [-0.017, -0.004]). The proportion bet was not different from the normative proportion on the first trial *(β*_*n*_ = 0.295, 95% CI = [0.219, 0.372]) but it deviated on the last trial *(β*_*n*_ = 0.179, 95% CI = [0.102, 0.256]). We did not find a downward trend in Game 2 of *O*_*n*_ *=* 3 (Figure 4E; slope = -0.003, 95% CI = [-0.010, 0.003]). The proportion bet was not different from normative proportion on both the first trial *(β*_*n*_ = 0.211,95% CI = [0.134, 0.287]) and the last trial *(β*_*n*_ = 0.177, 95% CI = [0.100, 0.253]).

A post hoc pairwise comparison with Bonferroni correction showed no significant difference in the average proportion bet for *O*_*n*_ = 3 between Game 1 and Game 2 (Δ = 0.018, SE = 0.016, *t* [901] = 1.11, *p* = 0.99) and in the average proportion bet for O_*n*_ = 4 between Game 1 and Game 2 (Δ = 0.022, SE = 0.016, *t* [901] = 1.34, *p* = 0.73), implying there was no significant effect of learning in Game 1 on performance in Game 2. People bet more in *O*_*n*_ = 4 than *O*_*n*_ = 3 in both game 1 (Δ = -0.048, SE = 0.016, *t* [901] = 2.94, *p* = 0.014) and game 2 (Δ = -0.044, SE = 0.016, *t* [901] = 2.77, *p* = 0.023), showing sensitivity to the different odds.

#### Experiment 1

*O*_*n*_ *-* 2: Gambles with *O*_*n*_ *-* 2 have an expected value of 0: they offer only risk and it is plausible that players should avoid them by setting *β*_*n*_ to 0. Nevertheless, players bet more than 0 tokens throughout the period of both game 1 *(β*_*n*_ for the first trial = 0.245, 95% CI = [0.190, 0.300]; *β*_n_ for the last trial = 0.096, 95% CI = [0.040, 0.152]) and game 2 *(β*_*n*_ for the first trial = 0.172, 95% CI = [0.117, 0.227]; for the last trial = 0.088, 95% CI = [0.033, 0.143]). When *O*_*n*_ *-* 2 players also bet less as the game progressed (slope for game 1 = -0.014, 95% CI = [-0.019, -0.008]; slope for game 2 = - 0.076, 95% CI = [-0.013, -0.002]).

### Property 2: [Experiment 1, Optimal Bet]

In Figure 5, we plot the mean proportion bet across trials for each player (marked as a black circle) for Experiment 1. We do not plot Experiment 2 in this format because of the difference in stopping rule. There are two panels AC for the plot of the mean proportion for *O*_*n*_ = 3 versus *O*_*n*_ = 2, separately for each of the games and two panels BD for the plot of the mean proportion for O_*n*_ = 3 versus *O*_*n*_ = 4. The expected rate of utility for each choice of fixed proportion *β* was computed using Monte Carlo simulation and plotted as a heatmap. A red square in each panel marks the normative proportion maximizing the rate of growth of

E[ logW ]. The ratio of normative proportions for *O*_*n*_ = 4 versus *O*_*n*_ = 3 was 0.333/0.25 = 1.33 the expected slope of the principal component line of the data. We estimated the ratio of player’s proportions in panels 4B and 4D by bootstrap simulation and found that the observed ratio was 1.371 (95% CI = [0.989, 1.562]) for game 1 and 1.164 (95% CI = [0.819, 1.428]) for game 2. We return to

#### Experiment 2

All the results for Experiment 2 are contained in the following Summary text box. We compared two conditions that differed only in stopping rule. There was no significant effect of stopping rule. Players did not attempt to maximize expected gain by betting all or most of their winnings on the last few turns.

**Summary: Experiment 2 [Effect OF Stopping Rule]**

- BET-IT-ALL [ EXPERIMENT 2, GAME 1 vs GAME 2 for *O*_*n*_*=3* and *O*_*n*_*=*4] *For O*_*n*_*=3* we test whether the proportion bet in the last triplet of Experiment 2, Game 2 is higher than the last trial of Experiment 2, Game 1? We found no significant effect (two-sample t-test, p = 0.416, t(27) = -0.82). *For O*_*n*_*=*4 we test whether the proportion bet in the last triplet of Experiment 2, Game 2 is higher than the last trial of Experiment 2, Game 1? We found no significant effect (two-sample t-test, p = 0.206 t(27) = 1.29).

## Discussion

A CAG is an abstract game based on probability and value, a relative of decision under risk, but with an emphasis on repeated play and accumulation. In a Cumulative Allocation Game (CAG) players have a store of wealth and – on every turn – risk a proportion of that wealth in order to gain more. CAGs are similar to decisions under risk in that both involve known probabilities and values on each turn. Three features that distinguish decision under risk from CAGs are, first, the players in a CAG choose how much or how little to risk on each turn; second, that a CAG with a constant hazard stopping rule. can go on indefinitely; and third, for many CAGs the player – with normative play – can expect exponential growth in wealth. An additional difference is that the order of trials of a decision under risk experiment is often randomized and players are given “one shot” instructions (Dillenberger, 2008; Guo, 2011): only one (or a small number) of the experimental trials will actually pay out money and the player will not learn which until the end of the experiment. The experimenter in a decision under risk experiment has done what he can to fix the expectations and strategies of the player. Every trial is effectively the first and only trial. In contrast CAGs are cumulative and the potential reward from future play is much larger than the immediate expected reward simply because the player is (probably) playing for higher stakes as the game progresses.

The design of the experiments (Figure 3) allowed us to test multiple hypotheses about the strategies adopted by human players and how they compare to normative strategies maximizing the expected rate of growth E[ logW ] or, in one condition maximum expected value.

In Game 1 and 2 of Experiment 1, players completed twelve trials (triplets) with each possible odds 2:1, 3:1,4:1 (a total of 36 trials, 12 trials per odds). Trials for different odds were interleaved. In Experiment 1 we compared performance in Game 1 and Game 2 to test whether players altered their strategies as they gained experience in the game. We concluded they did not.

In Experiment 1 players did not follow the normative strategy over the course of the game. However, if we consider only the initial trials of Game 1 and Game 2 we find that players’ *initial* bets for odds 3:1, 4:1 were not significantly different from normative (Eq. 3) but that, over the course of the game, players reduced their proportion bet in these conditions by a factor of 2. This failure reduced E[ logW ] by roughly 25%. We replicated this result in Experiment 1, Game 2.

### Why did initially normative subjects more away from the normative strategy?

To investigate possible reasons for this failure we examined what the players experienced in the CAGs. In Figure 6 we plot four time series for *O*_*n*_ *=* 3 for different proportions bet *β* The time series plotted are 500 turns long so that it easier to see the effect of changing *β* The green line is E[ logW ] for the choice of proportion *β\** that maximizes this growth rate, the normative strategy. The green time series is one realization of the experiment with the normative choice *b*-* Over many replications the resulting time series would be distributed symmetrically above and below the green line.

**Figure 6:**
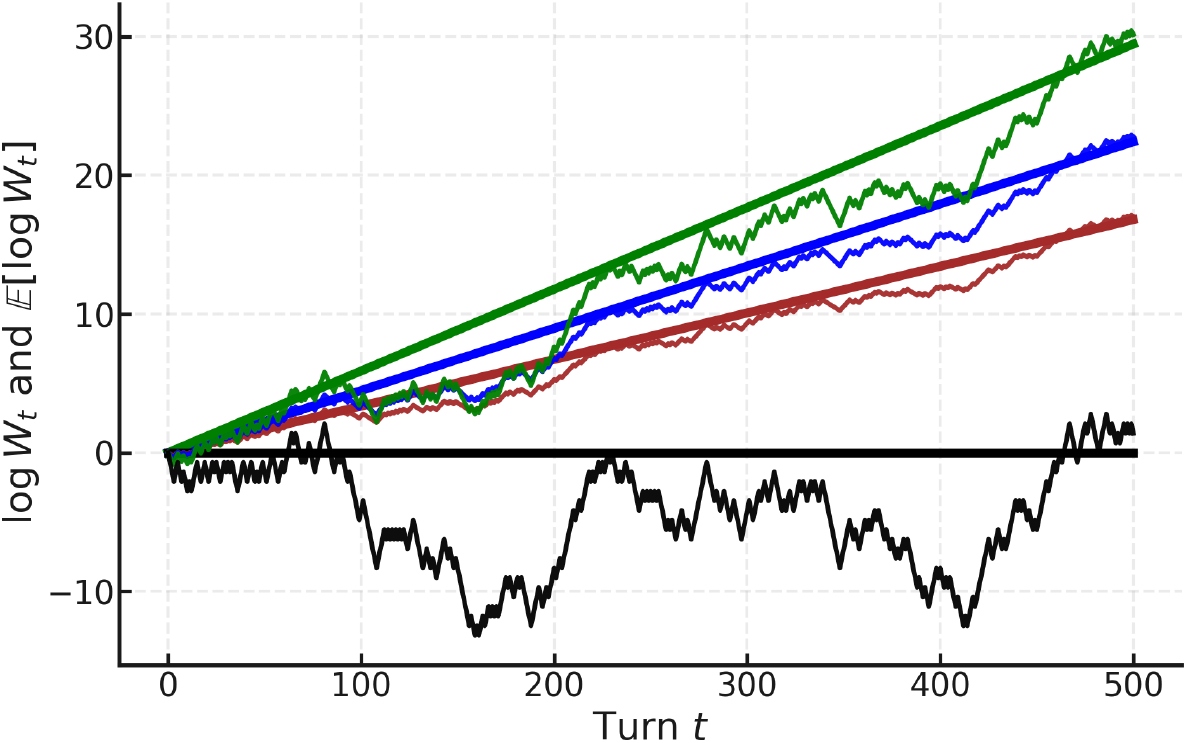
One simulated play of 500 trials of the the CAG with odds *O*_*n*_ *=* 3:1 for four different proportions bet. log [W_t_] is plotted against turn *t* for four different proportions bet. The straight lines are the color-coded expected means of many replications. The four values of *β* are 0.25 (green), 0.125 (blue), 0.0625 (red), and 0.5 (black). The value *β* maximizing the slope (rate of growth of log winnings) is β* = 0.25 (green), previously calculated (Eq. 3). We plot 500 trials to make the statistical patterns evident. This figure and the discussion surrounding it are taken from “The Kelly Criterion, Under the Hood” in Poundstone (2005). Following Poundstone, we used the same sequence of pseudo-random numbers for each time series to make them easier to compare.

The other lines and time series are simulations with *β* = *β*/2 and β = β*/3*, respectively. The player pursuing these strategies bets a fixed fraction of the normative proportion.

As we increase *β from β*/3 to β* the slope* E[ logW ] increases as does the volatility (“jaggedness”) of the time series.

Suppose we increase *β* further to *β = 2β\**. We add a single time series in black, with the slope E[ logW ] of the black line is flat and the black time series is extremely volatile. There are ranges where the black time series grows more rapidly than any of the others, but there are also frequent “collapses” where the time series returns to its starting point or below. Following such a collapse in the player’s fortunes the player can only slowly recover and any sustained growth is illusory. Another collapse will “surely” follow. Going beyond the normative bet proportion *b\** l eads to no increase in sustained growth E[ logW ].

The effect of increasing *β* is better seen if we compare just the black line and time series (β = 0.5) to the green line and time series in Figure 7 with 95% confidence regions (shaded green and grey). There are evident fluctuations in the green time series but the fluctuations are more and more unlikely to cancel out accumulated growth as the green line and the green confidence region trend upward away from 0. The confidence region for the black time series in contrast is subject to large downward fluctuations that preclude any sustained growth (Figure 7).

**Figure 7:**
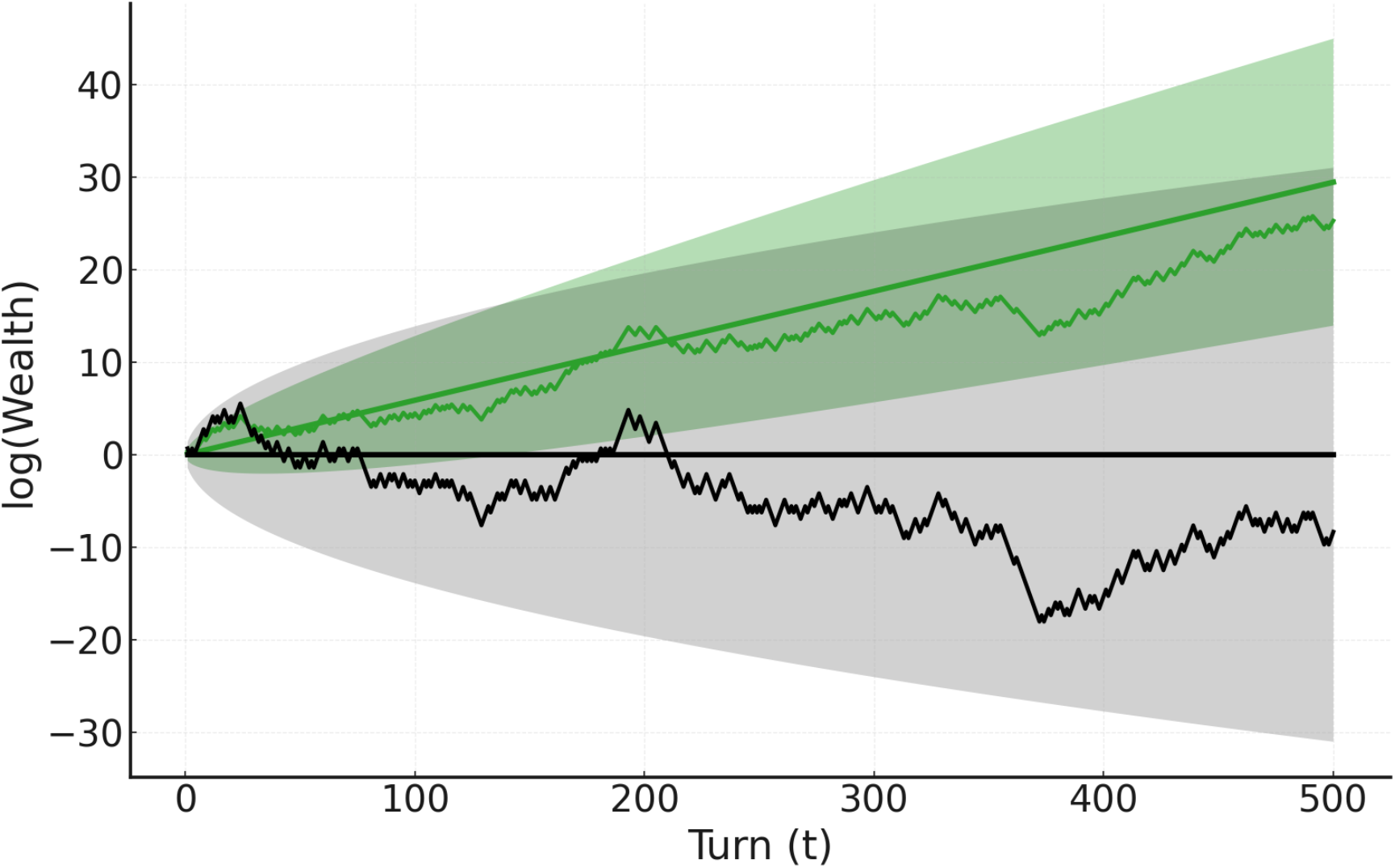
Time series of log [ W_t_ ], expected value of the time series (line) and a 95% confidence region for *β* = 0*.*25* (shaded green) and for *β =* 0.50 (shaded green). The grey region shows no trend while the green region does. The mean line of the green time series also trends upward and as the number of turns increases, the green series is less and less likely to return to lower values of E[ log W_**t**_ ]. We used the same pseudorandom number generator for both series to make them easier to compare.

Poundstone (2005) hypothesized that the behavior of human players is shaped by a tradeoff between E[ logW ] and volatility and our results are consistent with his hypothesis. As *β* increases up to the normative value *β\** we have linear growth in log [ W_**t**_ ] but also larger and larger fluctuations upward and downward that can wipe out the player’s wealth no matter how often he rebuilds it. In mathematical terms, *β\** is a singularity where the large fluctuations in the black series preclude any sustained growth in the expected mean (the black line). While *(\** is the proportion to bet that maximizes E[logW ] the player may find “the ride” a bit nerve-wracking.

The failure of proportionality in Experiment 1 is roughly a drift from the green line (*β\** = 0.25) and the corresponding time series to roughly the blue line and corresponding time series (β = 0.125). Why do players drift downward from normative play to subnormative?

By reducing the proportion bet to *β\**/2 or even *β\**/3, the player can still expect sustained, exponential growth but at a lower rate than would be possible with the normative strategy.

#### Loss aversion

We conjecture that this trend is not due to any feature unique to CAGs but is instead the consequence of a well-documented failing in human decision under risk: loss aversion (Kahneman & Tversky, 1979; Markowitz, 1952; Tversky & Kahneman, 1992) found that decision makers choosing between lotteries assigned more than twice as much weight (2.25 estimated experimentally) in calculating expected prospect value. This reweighting is captured in their prospect value function.

Return to Figure 6 and suppose that we apply a learning algorithm that corrects the log proportion bet by adding or subtracting a fixed percent feedback proportional to the absolute error with constant of proportionality ε > 0. This would result in a time series that would have no net tendency across trials to increase or decrease. If, however, excursions downward are (subjectively) weighted 2.25 times as much as excursions upward, then the corrected time series will trend downward at a rate determined by the correction percentage. This simple model would predict that loss aversion would lead to a reduction in proportion bet and a linear (in log) trend downward, the trend we observe. Moreover this reduction is arguably normative in that it represents a valid tradeoff between the volatility of the CAG and the rate of growth of wealth.

Simple economic games are a powerful tool when used to investigate the factors and computations that control human choice. Any model of human choice for one economic game, though, should make contact with other games, sharing the same factors and computations. If loss aversion in fact underlies the decreasing proportions bet then one aspect of classical decision under risk captured by Cumulative Prospect Theory (Tversky & Kahneman, 1992) is important in CAGs as well.

## Acknowledgments

YEW: Dean’s Undergraduate Research Fund, New York University, Twice awarded [April 2019, Nov 2019]; KO: Grant-Aid for JSPS Fellows [No. 17J07822)]; TMD funded by a Wellcome Career Development Award: [306332/Z/23/Z]; LTM: Humboldt Research Prize, Alexander von Humboldt Stiftung. We thank Lilly Li for help in executing the experiment, Puneet Bhargava for help in programming the experiment and members of the Vision & Decision Lab at NYU for discussion.

## Footnotes

1 We defined odds differently from Kelly (1956): our odds 2:1 is b = 1:1 in his notation.

